# PHYLOGENETIC ANALYSIS OF THE *rfb* LOCUS GENES OF THE GENUS *Leptospira* OF SEROGROUPS SERJOE, MINI AND HEBDOMADIS

**DOI:** 10.1101/2023.09.19.558452

**Authors:** Ruth Flávia Barros Setúbal, Jorge Estefano de Santana Souza, Maria Raquel Venturim Cosate, Tetsu Sakamoto

**Affiliations:** Bioinformatics Multidisciplinary Environment (BioME), Instituto Metrópole Digital (IMD), Universidade Federal do Rio Grande do Norte, Natal, RN, Brazil; MassBiologics, University of Massachusetts Medical School, Boston, USA

**Keywords:** Leptospirosis, *rfb* locus, Lipopolysaccharides, serological classification

## Abstract

Leptospirosis is a zoonosis of great impact on public health since it is considered a notifiable disease occurring mainly in tropical regions with poor sanitation and vulnerable socioeconomic conditions. It is caused by bacteria of the genus Leptospira and phylum Spirochaetes and contamination occurs through direct or indirect contact with the contaminating agent. In addition to taxonomic classification, which is performed through sequencing and the analysis of some marker genes, such as 16S rRNA and *secY*, they are usually classified based on their antigenic characteristics into serogroups and serovars. This kind of classification is largely applied in epidemiological studies and vaccine development. Despite its importance, few studies have been conducted to understand the evolutionary dynamics of the emergence or change of serology in this genus. In view of this, we applied phylogenetic methods in order to understand the evolutionary processes involving the serology of the genus. To this end, sequences of genes comprising the *rfb* locus from samples of serogroups Sejroe, Mini, and Hebdomadis (34 samples) were extracted and submitted to the phylogenetic pipeline, resulting in the inference of 75 maximum likelihood trees. Topology tests showed that most of the gene trees are significantly different from the species tree. We could depict the occurrence of lateral gene transfer between *L. borgpetersenii* and *L. kirschneri*; and *L. interrogans* and *L. weilli*. In this analysis, no evidence was found for the lateral gene transfer between samples of the Hardjo serovar of *L. interrogans* and *L. borgpetersenii*. Thus, it is also suggested that the occurrence of horizontal transfer of genes from the *rfb* locus between distinct species is less frequent than expected.

## 1. Introduction

Leptospirosis is a zoonosis with a major impact on public health, as it is considered a notifiable disease, occurring mainly in tropical regions with poor basic sanitation and vulnerable socioeconomic conditions. Characterized as a systemic infection, it is caused by pathogenic species of the genus *Leptospira* that infect and lodge in the renal system of reservoir animals, transmitting the disease to humans through direct or indirect contact with contaminated urine (de Brito et al., 2018; Mwachui et al., 2015).

The clinical manifestations can vary from asymptomatic to symptomatic, and the symptomatic form is often confused with other pathologies such as dengue and yellow fever. In milder cases of the disease, it is common for the patient to have yellow skin and eyes, as well as muscle pain. More severe cases include high fever, bleeding, respiratory failure, vomiting, diarrhea, dark urine, and jaundice (Rajapakse, 2022).

Bacteria of the genus *Leptospira* belong to the order Spirochaetales and family Leptospiraceae. They are characterized by long spiral forms, which have flagella coated on the outer membrane and are thus undetectable on the bacterial surface (Mwachui et al., 2015). These flagella are considered an important invasion mechanism, enabling the bacteria to move in liquid media and human tissues. Strictly aerobic and nutritionally fastidious, their growth is slow and can take up to two weeks, making it difficult to classify and identify strains in clinical samples (Mwachui et al., 2015; Vincent et al., 2019).

There are two types of classification for these bacteria: serological and taxonomic. Both are important for understanding diversity and evolution, as well as for developing strategies to prevent and treat leptospirosis (Khalil et al., 2021). Serological classification is based on the antigenic reaction of bacteria with specific antibodies. This classification is carried out using seroneutralization, agglutination, and ELISA techniques. The antigenic reaction allows the identification of the *Leptospira* species present in clinical samples, in which the *Leptospira* strains are classified into serovars (Latosinski et al., 2018). The division of serovars is based on the similarity of the structure of the lipopolysaccharide (LPS) carbohydrates present in the outer membrane. Some samples from different serovars can react to the same antigen, which is called a cross-reaction. Serovars that cross-react are further grouped into serogroups. The genus has around 24 serogroups and more than 250 serovars (Mejía et al., 2022). Meanwhile, the taxonomic classification is based on the analysis of the genetic material of these bacteria, including the sequencing of the complete genome or specific fragments. Genetic classification allows the identification of genetic divergences between *Leptospira* species, and the understanding of the evolution and diversification of these bacteria (Balassiano et al., 2017). The genus has 68 species identified so far and they are distributed in two major clades, one comprising the pathogenic species and other saprophytic species (Vincent et al., 2017).

The combination of serological and genetic classification has been useful for the development of more accurate and sensitive diagnostic strategies, as well as for the development of specific vaccines and therapies for leptospirosis, including the identification of prevalent serovars in different regions and the assessment of the geographical distribution of these bacteria (Balassiano et al., 2017; Mejía et al., 2022). However, there is still much to improve, and understanding the source of the genetic and serological diversity is fundamental to addressing more effective strategies for the prevention and control of leptospirosis. Recent studies showed a high association between the gene composition of the *rfb* locus and the serological classification (Medeiros et al., 2022; Nieves et al., 2022). In this sense, we aimed to analyze the evolutionary history of the genes comprising the *rfb* locus of the *Leptospira* genus in order to understand the evolutionary dynamics surrounding the serological classification of this genus. In this work, we proposed to carry out this analysis using genomic data from samples of Sejroe, Hebdomadis, and Mini (SHM) serogroups.

## 2. Material and Methods

### 2.1. Data source

The genomic data for all the samples used in this work (Table 1) was obtained from the NCBI database. From the available data, the FASTA files of the CDS of the annotated genes and the tables containing the genome annotations (feature table) were downloaded. Data on the serological classification of each sample was obtained from PubMLST (https://pubmlst.org; Jolley & Maiden, 2010). For samples that did not present serological data, we used the serological classification suggested by Medeiros et al. (2022), which is based on the gene profile of *rfb* locus.

**Table 1:**
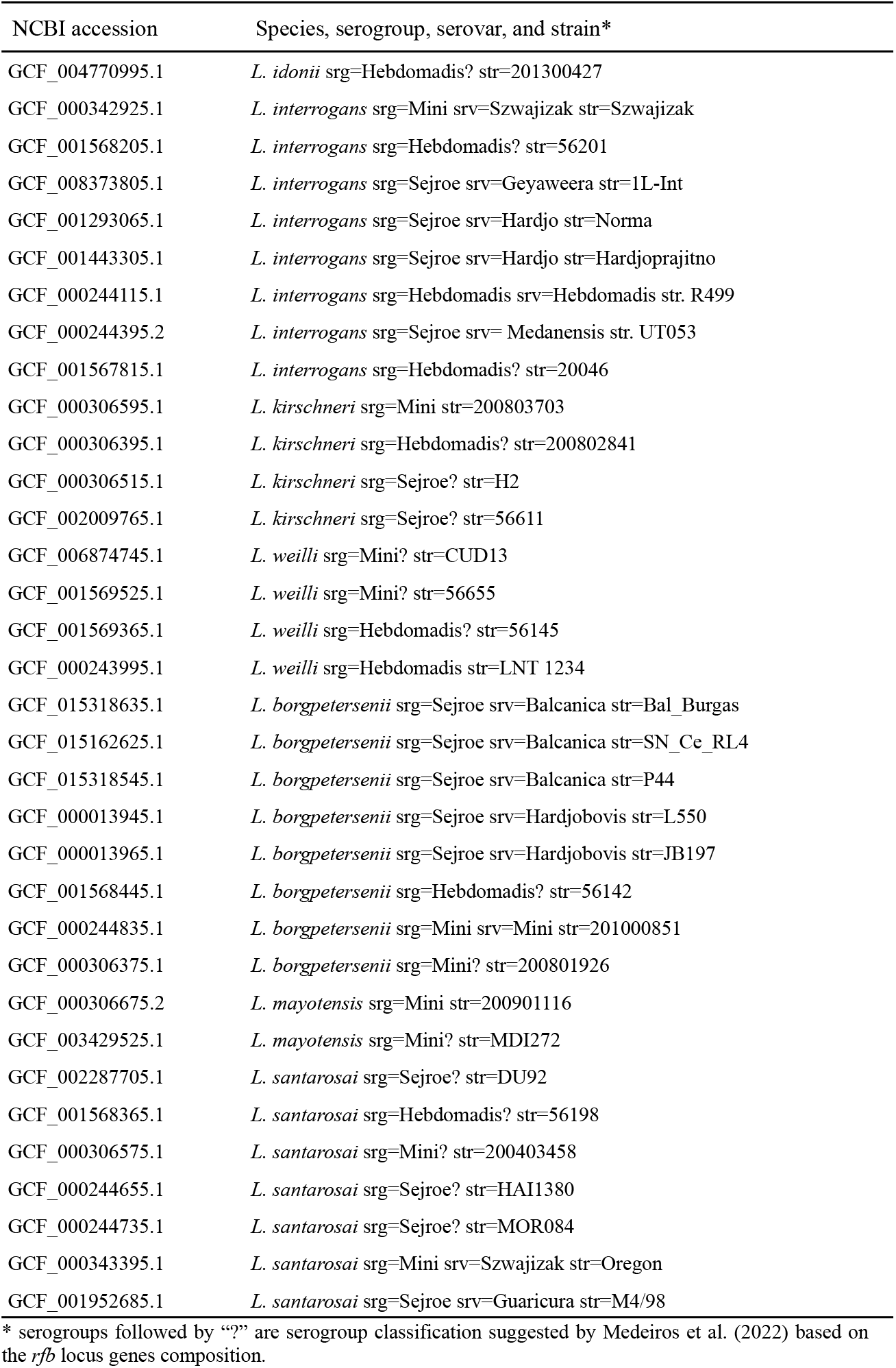
Samples used in this work.

### 2.2. Ortholog groups

Orthofinder 2 software (Emms & Kelly, 2019) was used to establish the groups of orthologs between the proteins. Before submission, the FASTA files of the CDS were translated into proteins using the seqkit program (Shen et al., 2016). The Orthofinder 2 pipeline was run using the default parameters.

### 2.3. Phylogenetic analysis

For a given orthologous group, we first extracted the CDS sequences of its members and translated them into amino acid sequences using the seqkit software (Shen et al., 2016). These sequences were aligned using the MUSCLE (Edgar, 2021). We then back-translated the alignment into nucleotide and submitted it to the trimAl (Capella-Gutiérrez et al., 2009), which identified and removed sites with low-quality alignments. All the alignments generated were inspected manually using the Mega software (Tamura et al., 2021). The resulting alignment was then submitted to the iqTree (Nguyen et al., 2015) to generate the maximum likelihood tree. To do this, parameters were used to select the evolutionary model and to carry out the bootstrap test with 1000 replications (-bb 1000). The trees generated were submitted to TaxOnTree (Sakamoto & Ortega, 2020) for annotation and visualized using Figtree (http://tree.bio.ed.ac.uk/software/figtree/).

### 2.4. Tree topology tests

To compare the topology of the maximum likelihood tree with other topologies, tree topology tests were carried out using the iqTree tool (Nguyen et al., 2015). The topology tests used in this analysis were: Bootstrap Proportion (BP), Kishino-Hasegawa test (KH, Kishino and Hasegawa, 1989), Shimodaira-Hasegawa test (SH, Shimodaira and Hasegawa, 1999), expected likelihood weights (c-ELW, Strimmer and Rambaut, 2002) and approximately unbiased test (Shimodaira, 2002). For the KH, SH, and AU tests, p-values were returned in which a tree is rejected if p < 0.05. The bootstrap tests and likelihood weights (BP and c-ELW) return weights that are not considered p-values and these add up to 1 in the trees tested.

## 3. Results

### 3.1. Evolutionary relationship of *Leptospira* strains of Sejroe, Mini, and Hebdomadis serogroups

After obtaining the genomic data of the 34 *Leptospira* samples of the Sejroe, Hebdomadis, and Mini (SHM) serogroups, their CDS sequences were translated into amino acid sequences and then submitted to the Orthofinder program to perform ortholog grouping. In this analysis, a total of 125,709 genes were grouped into 5,735 orthogroups. Among these orthogroups, the average orthogroup size was 21.9, and 1,738 orthogroups were single-copy. Along with the orthogroup analyses, the Orthofinder program also inferred a species tree based on the single-copy orthogroups (Figure 1). In this tree, after rooting the by the sample of *L. idonii*, all the samples that belong to the same species formed a monophyletic group. The phylogenetic analysis also showed two main clusters, one comprising *L. interrogans* and *L. kirschneri*, and the other comprising *L. borgpetersenii, L. mayotensis, L. weili*, and *L. santarosai*. The topology of the inferred species tree was also consistent with the topology of other *Leptospira* phylogenomics studies (Vincent et al. 2019).

**Figure 1:**
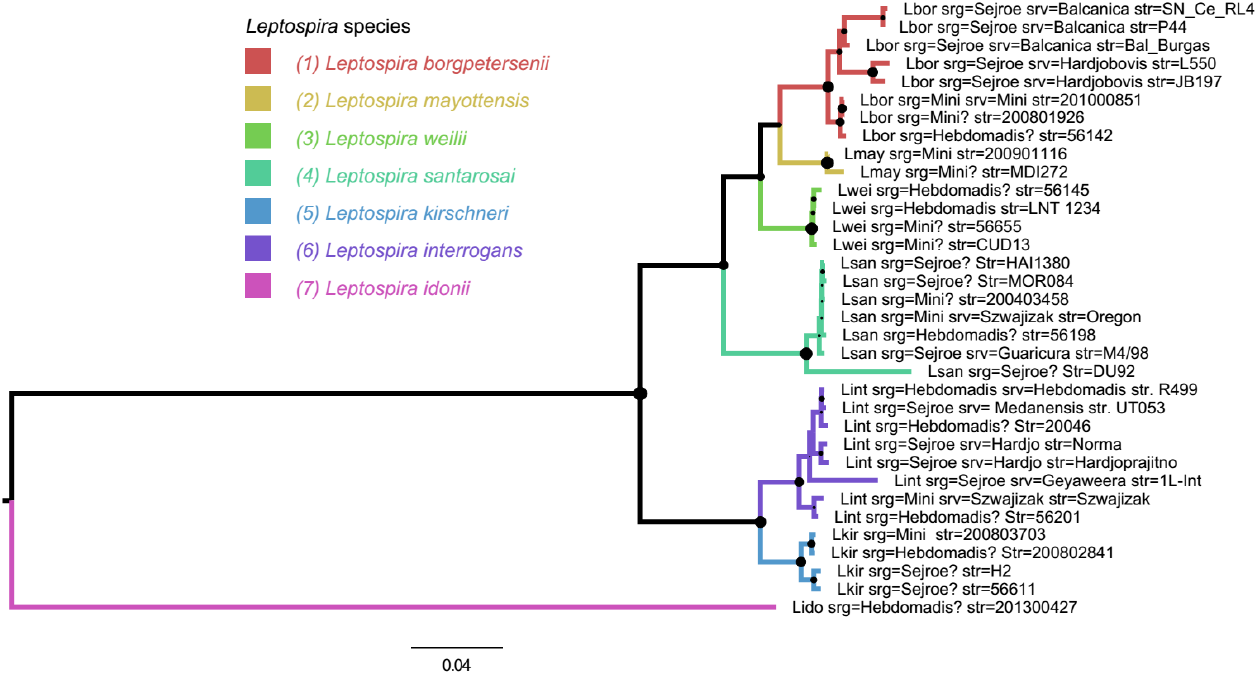
Species tree inferred by Orthofinder 2. Lbor: *L. borgpetersenii*; Lint: *L. interrogans*; Lmay: *L. mayotensis*; Lwei: *L. weilli*; Lsan: *L. santarosai*; Lkir: *L. kirchneri*; Lido: *L. idonii*; srg: serogroup; srv: serovar; str: strain.

### 3.2. Evolutionary analysis of the genes comprising the *rfb* locus of samples of SHM serogroups

Among the orthogroups, we identified the ones containing the genes that are part of the *rfb* locus. For this, we search for orthogroups containing the genes comprising the *rfb* locus of the *L. interrogans* str. Norma (srg Sejroe, srv Hardjo). In total, 75 orthogroups were identified and submitted to the phylogenetic pipeline. To make it easier to refer to these orthogroups, we have identified each of them by numbers from 1 to 75, following the order of occurrence of these genes in the *rfb* locus of *L. interrogans* str. Norma. It is noteworthy to mention that the gene structure of the *rfb* locus of pathogenic samples has a unique pattern. It starts with the MarR gene and ends with the DASS gene. Genes in the 3’ end of the *rfb* locus are conserved among all pathogenic samples and are composed of genes responsible for the O-antigen processing and synthesis through the Wzy-dependent pathway, and for the dTDP-rhamnose biosynthesis (*rfbC, rfbD, rfbB, rfbA*), which are involved in the assembly of LPS. The first two genes, MarR, and GDP-L-fucose synthase, also showed to be conserved among pathogenic samples. The gene composition in the remaining region of the *rfb* locus can differ considerably among samples of different serogroups, but samples of serogroups Sejroe, Mini, and Hebdomadis show similar composition. Considering the gene composition of the *rfb* locus of *L. interrogans* str. Norma, the first two genes (orthogroups 1 and 2) and the last 17 genes (orthogroups 59 to 75) are those conserved among pathogenic samples, and the remaining genes (orthogroups 3 to 58) are conserved among samples of SHM serogroups. Additionally, there is a set of four genes that are specific to samples of Sejroe serogroup (orthogroups 14 to 18). These genes are in a region called SHM rfb variable region (SVR), which can be used to distinguish samples between serogroups Sejroe, Hebdomadis, and Mini (Medeiros et al., 2022).

After inferring the maximum likelihood trees for each orthogroup, their topologies were tested in relation to the species tree. Of the 75 gene trees tested, only 12 (16.0%) showed a topology that was statistically consistent with the topology of the species tree. Seven of them are of genes conserved among the *Leptospira* samples, indicating that the evolutionary history of most of the genes in the *rfb* locus has been shaped by evolutionary events other than vertical transfer, such as horizontal gene transfer. We also performed a second topology test, this time testing monophyly between samples of the same species, and we found 31 trees being statistically consistent with the tested topology, indicating that interspecies lateral transfers could not be a frequent event.

By visually analyzing the tree topology of the maximum likelihood trees, we identified the occurrence of two main lateral transfer events. One involved the *L. borgpetersenii* (Lbor) and *L. kirschneri* (Lkir) species samples and the other involved the *L. weilli* (Lwei) and *L. interrogans* (Lint) species samples (Figure 2). To statistically verify these transfer events, tree topology tests were carried out, testing for monophyly between Lbor and Lkir, and between Lint and Lwei. The topology tests showed that monophyly between Lbor and Lkir, and between Lwei and Lint occurs in 44 and 28 orthogroups, respectively. In addition, orthogroups showing positive for both tests totaled 21. Most of these genes are those found only among SHM serogroups. Regarding the genes of the SVR region and specific to the Sejroe serogroup, the evolutionary history of the four genes showed the occurrence of lateral transfer between Lbor and Lkir. It was not possible to evidence lateral transfer between Lint and Lwei due to the lack of Lwei samples containing these genes.

**Figure 2:**
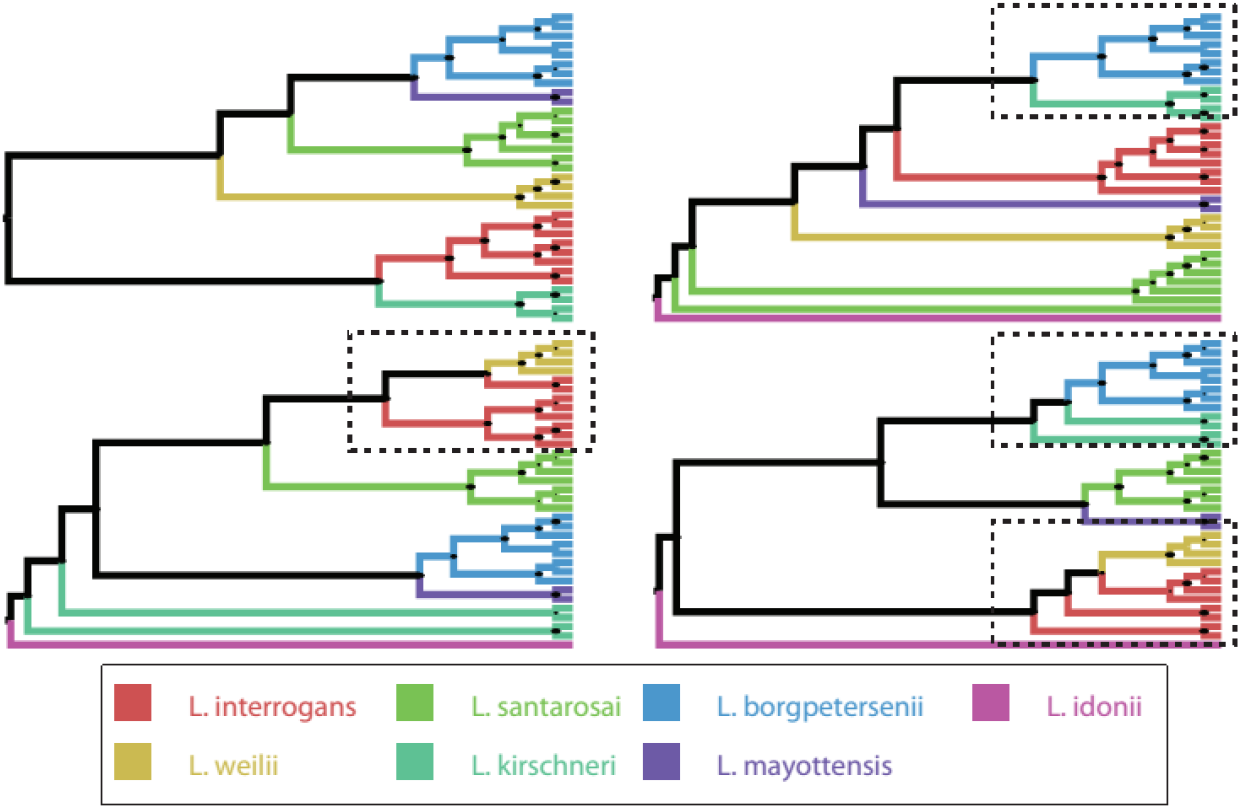
Different topologies of gene trees for the *rfb* locus inferred using the iqTree 2 program. Dashed rectangles indicate branches not found in the species tree, indicating possible lateral transfers.

## 4. Discussion

The first studies on the genetic factors responsible for determining the serogroup of the Leptospira genus were conducted with samples of the *L. interrogans* and *L. borgpetersenii* species that shared the same serological classification (srg Sejroe and srv Hardjo, Bulach et al., 2000; de la Peña-Moctezuma et al., 1999, de la Peña-Moctezuma et al., 2001). In these studies, it was found that the genes of the *rfb* locus of *L. interrogans* srv. Hardjo shared a high degree of similarity with the genes of the sample *L. borgpetersenii* srv. Hardjo. A similar analysis with another sample of the same species, but of a different serovar (*L. interrogans* serovar Copenhageni) resulted in a low similarity. This evidence provided support for the occurrence of lateral transfer events in this region closely associated with the serology of the genus. This is the best known and most cited example of an inter-species lateral transfer event within the community studying this pathogen.

With the growing number of *Leptospira* samples with genomic data deposited in public databases, Medeiros et al. (2022) were able to carry out a comparative genomic analysis of more than 700 Leptospira samples. In this work, they also found that it is possible to distinguish serological groups based on the constitution of the *rfb* locus. They also found that the Sejroe, Mini, and Hebdomadis serogroups shared a similar *rfb* locus gene profile, which explains the cross-reaction events that occur between them. In addition, it was found that there is a small region of the *rfb* locus, called the SVR (Sejroe/Mini/Hebdomadis rfb variable region) by the authors, which allows samples to be classified between these three serogroups according to their gene constitution. Other studies have also found an association between the *rfb* locus and serological classification in other species and serogroups (Nieves et al., 2022; Fouts et al., 2016).

Mejía et al. (2022) also verified the importance of recombination events in this genus throughout its evolution. Their analysis found that approximately 22% of the core genes had undergone a recombination event. This study also involved the genes responsible for LPS biosynthesis. However, their analysis only focused on the genes that are responsible for the synthesis of the Lipid-A portion, which is the portion that anchors the LPS to its outer membrane and which is outside the *rfb* locus. In this work, we focused our studies on the genes in the *rfb* locus that are responsible for synthesizing the O-antigen portion of an LPS. The O-antigen is the portion of the LPS that is exposed to the environment and determines the serological diversity between samples.

The phylogenetic analyses carried out in this study allowed us to contemplate some of the evolutionary events that permeate the serological classification of the *Leptospira* genus. Preliminary analyses showed that the *L. kirschneri* and *L. weilli* species had received some genes responsible for synthesizing the O-antigen structure of LPS through interspecies lateral transfer and that they had undergone changes in their serology. Despite this evidence of lateral transfer between different species, we also found that this event may not be as frequent as previously thought. We also did not find evidence of a recent event of lateral gene transfer between the samples of serovar Hardjo of *L. interrogans* and *L. borgpetersenii*, suggesting that the transfer event that originated the serovar Hardjo in one of these species may occurred longer ago. There is also a suggestion that there could be other genes outside the *rfb* locus that could be involved in the manifestation of the serovar Hardjo phenotype.

